# An exhaustive analysis of single amino acid variants in helical transmembrane proteins

**DOI:** 10.1101/2019.12.18.881318

**Authors:** Oscar Llorian-Salvador, Michael Bernhofer, Yannick Mahlich, Burkhard Rost

**Affiliations:** TUM, Department of Informatics, Bioinformatics & Computational Biology - i12, Boltzmannstr. 3, 85748 Garching/Munich, Germany; Department of Biochemistry and Microbiology, Rutgers University, 76 Lipman Dr, New Brunswick, NJ, 08901, USA; Institute of Advanced Study (TUM-IAS), Lichtenbergstr. 2a, 85748, Garching/Munich, Germany; Institute for Food and Plant Sciences WZW – Weihenstephan, Alte Akademie 8, Freising, Germany

**Keywords:** helical transmembrane proteins, single amino acid variants, functional features prediction, human proteome, topology prediction

## Abstract

Single nucleotide variants (SNVs) have been widely studied in the past due to being the main source of human genetic variation. Less is known about the effect of single amino acid variants (SAVs) due to the immense resources required for comprehensive experimental studies. In contrast, in silico methods predicting the effects of sequence variants upon molecular function and upon the organism are readily available and have contributed unexpected suggestions, e.g. that SAVs common to a human population (shared by >5% of the population) have, on average, more significant impact on the molecular function of proteins than do rare SAVs (shared by <1% of the population). Here, we investigated the impact of variants in a human population upon helical transmembrane proteins (TMPs). Three main results stood out. Firstly, common SAVs, on average, have stronger effects than rare SAVs for TMPs, and are enriched, in particular, in the membrane helices. Secondly, proteins with seven transmembrane helices (7TM, including GPCRs, i.e. G protein-coupled receptors) are depleted of SAVs in comparison to other proteins, possibly due to increased evolutionary constraints in these important proteins. Thirdly, rare SAVs with strong effect are significantly absent (over common SAVs) in signal peptide regions.

## Introduction

Single amino acid variants (SAVs) have been found to be relevant for the molecular function of proteins [1]. The impact of different SAVs has been predicted *in silico* showing that, within human, more common SAVs (observed in more than 5% of the population) tend to have a larger impact on protein function than the rare ones (observed in less than 1% of the population [1]. The effect of common SAVs (gain or loss of functionality) on the survival of the species remains, at least partially, unknown.

It was critical to continue the previous research on the functional impact of common SAVs to have a better understanding of their effect on the survival of species. A more thorough analysis of the effect of common and rare SAVs may lead to a significant connection between these micro-molecular variations and their phenotypic impact on an organism.

A group of proteins with a different proportion of variants, such as an enrichment in common SAVs associated with strong effect, can show locations or specific functionalities where a higher variability helps the whole species. It can also show locations or functionalities with less variability, indicating a benefit only for individuals. In addition, this analysis can be extended not only to groups of proteins but regions of proteins: is it possible to find an enrichment in common SAVs in a certain type of structure?

Here, we analyzed the proportion of common and rare human SAVs in helical transmembrane proteins (for referred to as TMPs; note that a very small fraction of all human transmembrane proteins cross the membrane with beta-strands). About 24% of all human proteins have at least one transmembrane helix (TMH), and about 13% have two ore more TMHs [2]. A separate analysis of SAVs in globular-water soluble and TMPs is supported by the functional differences of these types of proteins, in particular by the many roles in the cell of TMPs, e.g. for regulation, signalling and transport across membranes [3–5]. We largely refuted our initial hypothesis, namely that a substantial fraction of the signal why common SAVs have, on average, more effect than rare SAVs although we observed an over-representation of common effect SAVs in TMPs.

## Materials & Methods

First, we analyse common SAVs with strong effect on protein function in a specific group of proteins: TMPs. Multiple comparisons are used to show differences in functionality between common and rare SAVs in all proteins and only in TMPs.

Second, we study the SAV distribution within TMPs: is it possible that certain regions, for instance transmembrane helices (TMHs), are enriched for a specific type of SAV?

### Starting dataset

We took all SAVs in the human population from a previous study [1] based upon a raw dataset with all proteins and SAVs reported by the Exome Aggregation Consortium (ExAC) [6] collecting 60,706 human exomes. The dataset contained 10,474,468 SAVs. For about 73% of these, namely for 7,599,572, we could find sequences in the UniProt repository [7]. For all of those we predicted their effect upon molecular function using the machine learning based method SNAP2 [8]. Transmembrane helices (TMHs) were identified through the method TMSEG [2], also used to distinguish between globular-soluble and transmembrane proteins (TMPs).

For each SAV, the dataset also contained the following information: allele frequency, location within the protein, mutation description, type of mutation and the expected effect of the mutation. Allele frequency was used to divide SAVs according to their abundancy in the human population (here proxied by 60,706 individuals) into three types: **<u>common SAV</u>**: observed in over 5% of the population, **<u>rare SAV</u>**: observed in less than 1% of the population, and intermediate all in between (5%≥i≥1%) which were ignored in this study. Predicted SAVs were classified as predicted with **<u>strong effect</u>** for high SNAP2 scores above 0 (probability gradually increasing for more positive scores), and to predicted to be **<u>neutral</u>** for low SNAP2 scores below 0 (probabilty gradually increasing for more negative scores).

### Dataset refining

The starting dataset contains 70,339 proteins with an available transmembrane prediction and at least one SAV. Since there are about 20,000 different proteins in the human exome, it is possible that there are multiple isoforms of the same proteins and small peptides that needed to be removed from the analysis.

To this purpose, we filtered the dataset by mapping the potential isoforms to their canonical proteins: we mapped SAVs occurring in isoforms to the canonical protein sequences by retrieving the respective protein sequences from UniProtKB and aligning them with Kalign [9]. Then we calculated the positions of all SAVs according to those alignments, ignoring regions that differed in their amino acids between the isoforms and canonical sequences.

In addition, proteins shorter than 50 amino acids were also discarded since they are, most likely, not complete TMPs. Therefore, the total number of proteins in the dataset with available topology prediction and SAVs was reduced to 16,644 (Table 1).

**Table 1.**
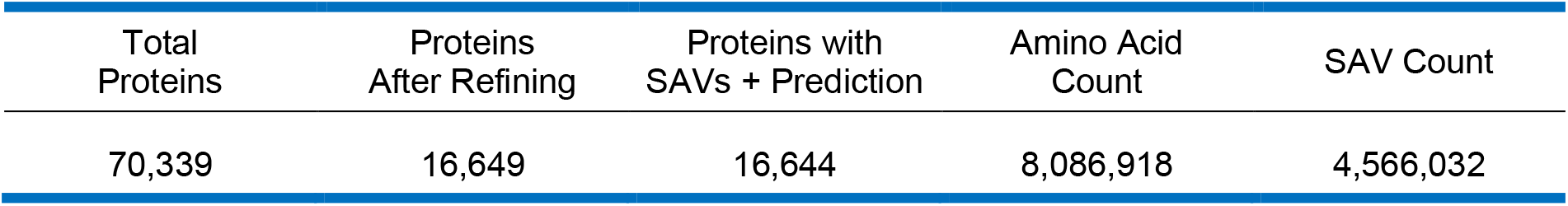
Protein count during the pre-processing step of the dataset. Last two columns show the amino acid and variant counts for the pre-processed dataset (16,644 proteins).

| Total Proteins | Proteins After Refining | Proteins with SAVs + Prediction | Amino Acid Count | SAV Count |
| --- | --- | --- | --- | --- |
| 70,339 | 16,649 | 16,644 | 8,086,918 | 4,566,032 |

### Validating the dataset

To validate the current dataset with the results obtained by the prior study [1], a comparison is made between common SAVs and rare SAVs with strong effect on protein function. In order to achieve results as similar as possible, the same parameter values are used:

- To define which SAVs are common and rare, we used the same allele frequency thresholds: SAVs with an allele frequency of 0.05 or higher are classified as common, SAVs with a frequency of 0.01 or lower are classified as rare. SAVs with a frequency between 0.01 and 0.05 are classified as uncommon, and will not be used in order to have two clearly separated groups.
- To define which SAVs have a significant effect on protein function, we used the same thresholds for the SNAP2 prediction score used in the previous work: SAVs with a SNAP2 score higher than 0 are likely to have a more significant effect on protein function than SAVs with a SNAP2 score below 0.

### Filtering the TMP dataset

Once the dataset has been validated, we continued the study with the proteins that have been predicted to be TMPs using TMSEG. The number of TMPs in the current dataset is 4,527 (27.2% of all proteins).

In a more thorough look at this TMP dataset, we compare the proportion of amino acids and variants to find possible enrichments. In addition, we repeat this comparison for different TMP types (single-pass, multi-pass and 7-pass TMPs), paying special attention to 7-pass TMPs (multi-pass TMPs that go through the membrane 7 times). This last group has special biological relevance, being the G protein-coupled receptors (GPCRs) part of this group [10–13].

### SAV analysis on the TMP dataset

Once the TMP group is examined, we investigate possible SAV enrichments within the protein types and regions. To find these enrichments, we extract the number of common and rare SAVs in the whole dataset and within each TMP type. Further, we compare the same groups regarding SAVs that have been predicted to have a significant effect on protein function.

However, the thresholds used are not completely binary: a SAV with a predicted SNAP2 score of 10 may still have a near-neutral effect on protein function; in the end, the threshold choice was decided arbitrarily. Likewise, the allele frequency thresholds used to define which SAVs are common and which ones are rare were chosen arbitrarily, although an intermediate group (uncommon variants) was defined so common and rare variants are clearly separated.

On the other hand, we analyse the effect of different SNAP2 score thresholds: with the use of several graphs, we show the proportion of SAVs that are predicted to have an effect on protein function, for all threshold values from −100 to 100 with a step of 1.

## Results and Discussion

Here we will discuss the analysis of Single amino acid variants (SAVs) in TMPs. First we validated the results with the ones from the previous work [1]. Then, we compared the effect on protein function and frequency of the SAVs in different protein groups, focusing on TMPs. Last of all, we will discuss how SAVs are distributed in the different TMP regions.

### Dataset validation

The results obtained by the prior research [1] show an enrichment in common variants associated with a strong effect on protein function (61% of the common variants had a significant effect), compared to the rare variants associated with a strong effect (50% of the rare variants had a significant effect).

In this study, a similar enrichment has been found: 65.5% of the common variants are predicted to have a significant effect on protein function, and 51.3% of the rare variants have a strong effect (Table 2). Even though the proportions are not the same, it was possible to confirm the enrichment of common variants associated with strong effect.

**Table 2.** Number of proteins, amino acids, and SAV distribution for three groups of proteins: all proteins, helical TMPs and other (soluble) proteins.

|  | Protein number | Amino Acids | SAVs | Common, strong effect SAVs | Rare, strong effect SAVs |
| --- | --- | --- | --- | --- | --- |
| All Proteins | 16,644 | 8,086,918 | 4,566,032 | 65.5 ± 0.6% | 51.31 ± 0.02% |
| Helical TMPs | 4,527 | 27.127% | 28.04 ± 0.02% | 62.2 ± 0.9% | 48.59 ± 0.06% |
| Soluble Proteins | 12,117 | 72.873% | 71.96 ± 0.02% | 67.1 ± 0.7% | 52.37 ± 0.03% |

### SAV distribution: helical TMPs and other proteins

Using the predictions made by TMSEG, two different groups of proteins were distinguished: TMPs and non-TMPs (referred to as soluble proteins). For these two groups, we have analysed the distribution of amino acids and SAVs, and the proportion of common and rare SAVs with a strong effect on protein function.

First, by comparing the proportion of amino acids and variants for helical TMPs, it is possible to see a small enrichment in variants in this group. In addition, a similar enrichment found in the complete dataset is present in both groups: a significant proportion of common variants have a strong effect on protein function.

Looking at the TMPs group, the proportion of all SAVs with a strong effect associated is lower compared to the complete set of proteins, regardless of the SAV frequency (TMP dataset: 62.2 ± 0.9% for common variants, and 48.59 ± 0.06% for rare variants; complete dataset: 65.5 ± 0.6% for common variants, and 51.31% ± 0.02% for rare variants).

On the other hand, looking at the soluble proteins group, there are significantly more common SAVs with effect compared to the TMP dataset (67.1 ± 0.7% versus 62.2 ± 0.9%). This proportion of common SAVs with associated effect is also higher than the same proportion in the complete dataset (65.5 ± 0.6%). In addition, the same can be observed when comparing rare SAVs between the complete dataset, soluble proteins and TMPs.

### Analysing SAVs on the TMP dataset

There are 4,527 TMPs in the complete dataset (which constitute a 27.2% of all proteins in the dataset), of which 42.3% are single-pass TMPs and 57.7% of them are multi-pass TMPs. Out of all TMPs, 16.5% of them are 7-pass TMPs (Table 3).

**Table 3.** Protein distribution for the different TMP types in the dataset. The Multi-pass TMP column also includes 7-pass TMPs.

|  | TMPs | Single-pass TMPs | Multi-pass TMPs | 7-pass TMPs |
| --- | --- | --- | --- | --- |
| Proteins | 4,527 | 1917 (42.3%) | 2610 (57.7%) | 749 (16.5%) |

There are 1,280,518 TMP SAVs (28.04% of all variants), which compared to the protein proportion (27.199%), indicates that there may be a variant enrichment in TMPs. In addition, there are 2,193,742 TMP amino acids (27.127% of all amino acids), which also shows a possible enrichment of SAVs in TMPs at the amino acidic level.

### Analysing SAVs on the TMP dataset: TMP types

A comparison between the proportion of amino acids and SAVs can show potential variant enrichments, meaning that for each amino acid there are more variants.

For all three types of TMPs, the proportion of amino acids and variants is similar, with only small differences: the proportion of single-pass TMP SAVs is a little higher than the proportion of amino acids (44.79% and 45.54%, respectively; Table 4). On the other hand, for multi-pass TMPs and 7-pass TMPs, this proportion is slightly lower (55.210% vs 54.46% and 12.659% vs 11.65%, respectively; Table 4).

**Table 4.**
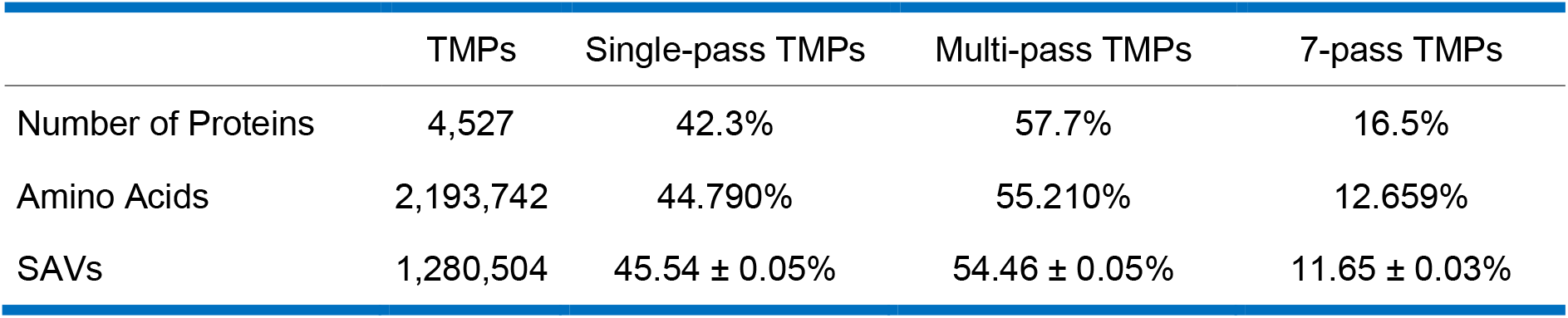
Number of proteins, Amino acids and SAVs for the different types of TMPs in the dataset.

It has been reported that the proportion of common and rare SAVs are very uneven in the human proteome: the majority of variants are rare (99%), while only 0.5% of the variants are considered common [6, 14]. Taking this in regard, the percentages of common and rare variants with a strong effect associated were obtained based on the variant frequency (Table 5 and other tables further on), so common variants are compared to common variants, and rare variants to rare variants. For instance, there are 2,053 variants in single-pass TMPs that are common and show a strong effect on protein function; these 2,053 variants constitute the 61% of all common SAVs (3,345 SAVs) in single-pass TMPs. We can observe that multi-pass TMPs have a higher proportion of common, strong effect SAVs (Table 5). This is especially noticeable when looking at 7-pass TMPs, which are clearly enriched for these variants.

**Table 5.** Distribution of common and rare SAVs with a strong effect associated in the different TMP types.

|  | All TMPs | Single-pass TMPs | Multi-pass TMPs | 7-pass TMPs |
| --- | --- | --- | --- | --- |
| Common, strong effect SAVs | 62.2 ± 0.9% | 61 ± 1% | 63 ± 1% | 68 ± 2% |
| Rare, strong effect SAVs | 48.59 ± 0.05% | 48.19 ± 0.09% | 48.92 ± 0.08% | 50.1 ± 0.2% |

When looking at different SNAP2 threshold, common variants always tend to have a higher effect on protein function than rare variants (Fig. 1). This difference is distinguishable using almost any threshold value, being clearer with non-extreme threshold values. In addition, confirming what we could already see (Table 5), 7-pass TMPs clearly contain the variants with the strongest effect among all TMP types, while the rest show a weaker, similar effect.

**Figure 1.**
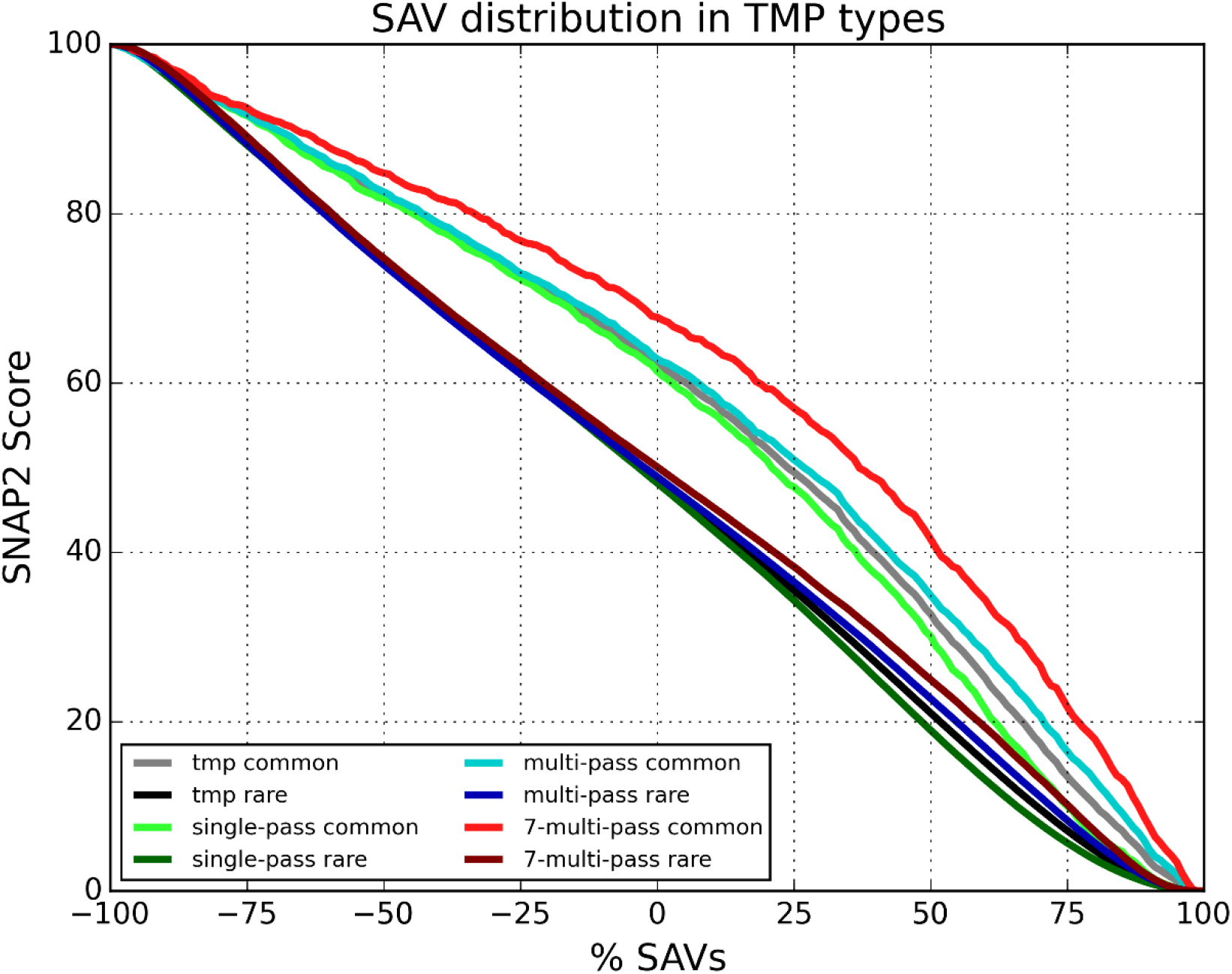
Common and rare variants predicted to have an effect on TMP function. All TMP types are presented. Y axis: Percentage of SAVs that have been predicted to have an effect. X axis: SNAP2 threshold used to predict that a variant has effect on protein function. Light colours stand for common variants, and darker colours for rare variants.

Regarding the rare variant distributions, they are all below their common variant counterparts, with small differences between them. At lower threshold values they are barely distinguishable (negative thresholds). On the other hand, at higher threshold values, it is possible to see that rare SAVs from 7-pass TMPs are slightly better predicted to have an effect on protein function, while rare SAVs from single-pass TMPs have the smallest predicted proportion overall.

Finally, at high threshold values (60+), the difference in the proportion of strong-effect SAV prediction between common and rare variants decreases, reaching a point where all groups of variants are separated by a similar percentage of SAVs predicted with effect (threshold value of 70), with common variants still having a higher proportion of SAVs predicted with effect.

### A deeper look into 7-pass TMPs

Given the special biological relevance of this group of proteins, here we analyse in detail their SAV distribution along their regions, frequency and effect.

If we compare the proportion of amino acids and variants for each region, a clear absence of variants in the TMH region is found. Since these proteins usually have a very specific function [10–13], variants would be more easily discarded due to the risk of modifying the protein function, or changing their structure along the membrane. This would therefore decrease the proportion of variants in this region.

Consequently, the proportion of SAVs in non-TMH regions is significantly higher. The signal peptide region, does not have a clear enrichment or absence in terms of variants.

In order to analyse the frequency and effect of SAVs in 7-pass TMPs, we have to take into account, again, the irregular distribution of common and rare variants in the human proteome [6, 14]. It is, therefore, necessary to split the SAVs using the frequency when looking at their predicted effect on protein function as well: otherwise such comparison would be biased towards rare variants.

We saw that signal peptide regions constitute 0.085% of the amino acids in 7-pass TMPs (Table 6). To take this uneven distribution into account, we have calculated SAV-amino acid index which weighs the region size by considering the number of variants per amino acid (Table 7, last two rows). We show that there is little difference in common, strong effect variants between TMH and non-TMH regions: non-TMH index is only slightly higher, pointing out that this region has a few more variants that meet these criteria.

**Table 6.** Distribution of amino acids and SAVs in the 7-pass TMP regions of the dataset. Non-TMH regions are excluding the signal peptides.

|  | 7-pass TMPs | TMH Region | Non-TMH Region | Signal Peptide Region |
| --- | --- | --- | --- | --- |
| Amino acids | 277,707 | 41.406% | 58.510% | 0.085% |
| SAVs | 149,134 | 36.9 ± 0.2% | 63.0 ± 0.2% | 0.070 ± 0.007% |

**Table 7.** Distribution of common and rare SAVs with a strong effect associated in the 7-pass TMP regions. Last two rows show the variant / amino acid ratio (called index) to consider the distribution of amino acids among the TMP types.

|  | All 7-pass<br>TMPs<br>Regions | 7-pass TMH<br>Region | 7-pass Non-TMH<br>Region | 7-pass Signal<br>Peptide Region |
| --- | --- | --- | --- | --- |
| Common, strong effect SAVs | 68 ± 2% | 67 ± 3% | 68 ± 3% | 0% |
| Rare, strong effect SAVs | 50.1 ± 0.2% | 54.6 ± 0.3% | 47.4 ± 0.2% | 19 ± 4% |
| Common, strong effect SAV index | 3.6e-03 | 3.6e-03 | 3.7e-03 | 0 |
| Rare, strong effect SAV index | 0.265 | 0.257 | 0.270 | 0.081 |

Given that there are only 749 out of 4,527 TMPs that are 7-pass TMPs (16.5% from Table 3), that signal peptide regions constitute only 0.085% of the amino acids in these proteins (Table 6), and the reported irregular distribution that common and rare variants usually have [6, 14], it is not very surprising that there are no common variants with a strong effect associated in signal peptide regions from 7-pass TMPs. Moreover, there are only 20 of their rare counterparts, resulting in a proportion of 19 ± 4% and an index of 0.081. Since this number of SAVs is too low to draw any conclusion (reflected in a high standard deviation), further analyses of these variants are recommended.

When using different SNAP2 thresholds to analyse common and rare SAVs on 7-pass TMPs, it is confirmed that there are no common variants in the signal peptide region for 7-pass TMPs (observed in Table 7) (Fig. 2). In addition, the low resolution for rare SAVs in the signal peptide regions confirms that there are not enough rare variants in this region to draw conclusive results (Table 7 for a SNAP2 threshold of 0).

**Figure 2.**
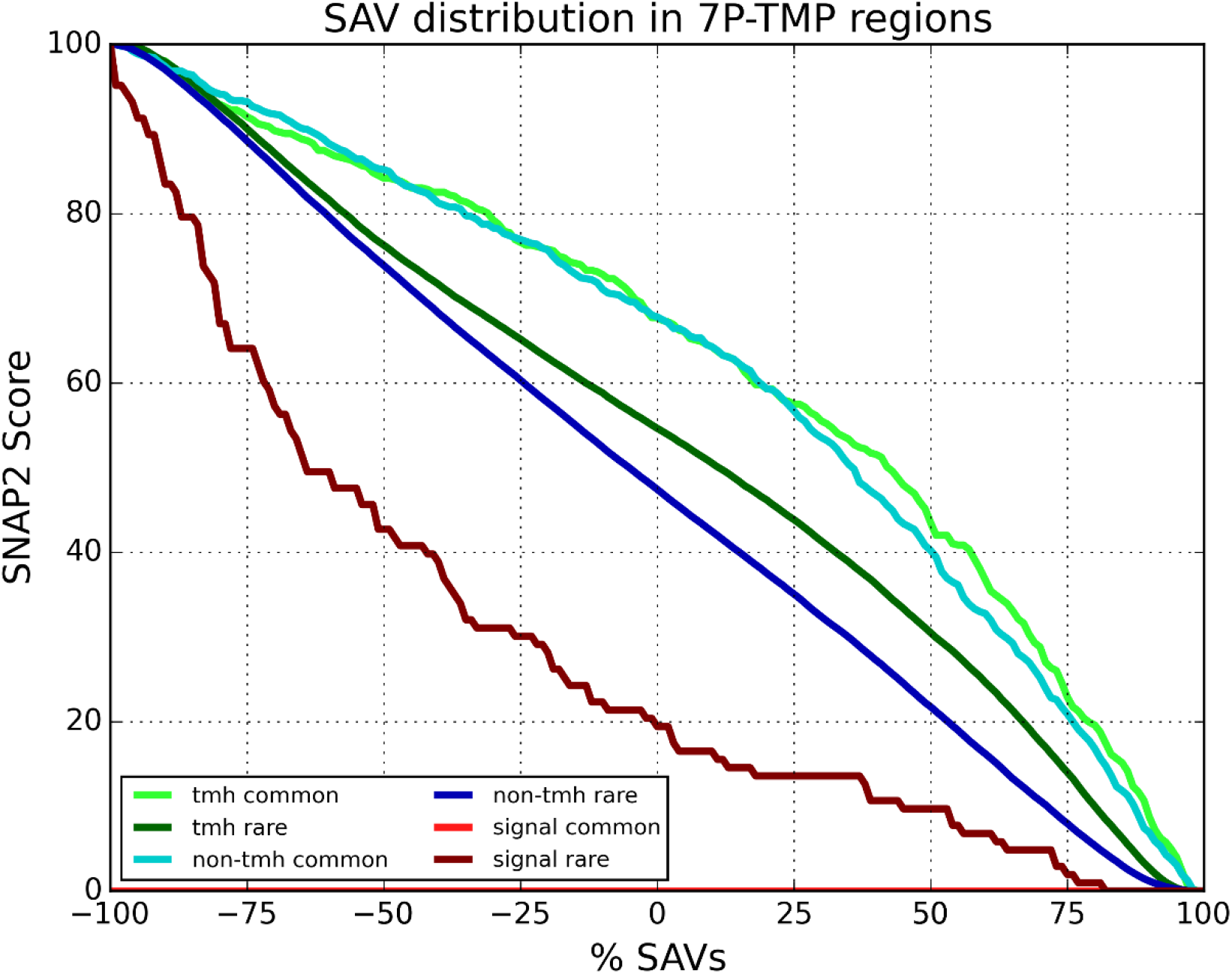
Common and rare variants predicted to have an effect on 7-pass TMP function. Variants are grouped by the protein region they appear in. Y axis: Percentage of SAVs that have been predicted to have an effect. X axis: SNAP2 threshold used to predict that a variant has effect on protein function. Light colours stand for common variants, and darker colours for rare variants.

In addition, all common variants have a higher effect on protein function for almost any prediction threshold when compared to rare variants (reflected in Fig. 1 and Fig. 2). Regarding TMH and non-TMH regions, both show a very similar proportion of common SAVs with any degree of effect on protein function, although the proportion may be slightly higher for TMHs when higher threshold values are used (above 25).

### Analysing SAVs on the TMP dataset: TMP regions

SAV enrichments can not only be found in specific types of TMPs, but also in specific TMP regions, which gives us a different, interesting perspective.

To test whether there are enrichments that may depend on TMP regions, we investigate the distribution of amino acids and SAVs classified by the TMP region they belong to.

Non-TMH regions constitute most of the SAVs (81% of the TMP amino acids and contain 82% of the TMP variants; Table 8). This uneven distribution needs to be accounted for when comparing results between TMP regions, for which a SAV / amino acid index has been already introduced (Table 7). Lastly, TMH and signal peptide regions have a slight lower proportion of variants compared to non-TMH regions.

**Table 8.**
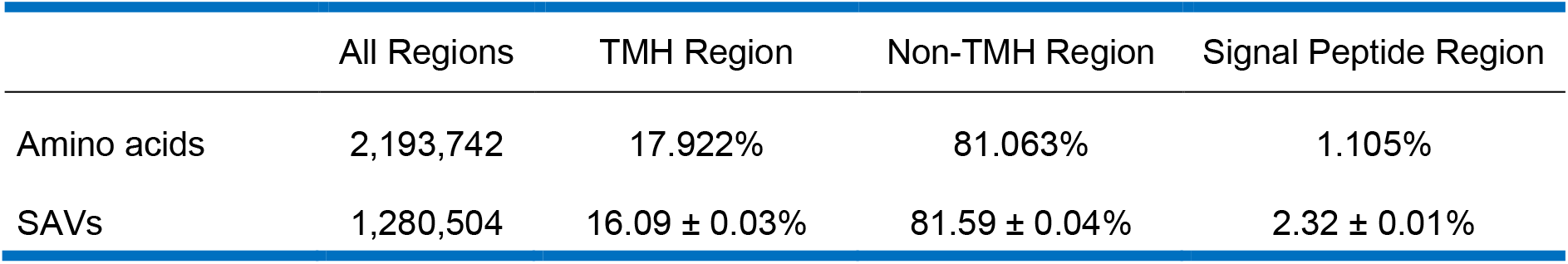
Amino acid and SAV distribution for the TMP regions: TMHs, non-TMHs and signal peptides. Non-TMH regions exclude signal peptides as well.

|  | All Regions | TMH Region | Non-TMH Region | Signal Peptide Region |
| --- | --- | --- | --- | --- |
| Amino acids | 2,193,742 | 17.922% | 81.063% | 1.105% |
| SAVs | 1,280,504 | 16.09 ± 0.03% | 81.59 ± 0.04% | 2.32 ± 0.01% |

Regarding the number of common and rare variants in TMPs in general, we see that 1,268,028 (99.03%) of the SAVs are rare, while only 7,309 (0.571%) of them were predicted as common (Table 9). It can be confirmed that it is possible to find approximately the same proportions when comparing the human TMPs and the complete human proteome [6, 14].

**Table 9.** Distribution of SAVs that are common and rare for each TMP region. Since there are uncommon variants that are not considered (allele frequency between 0.01 and 0.05), both percentages do not add to 100%.

|  | All regions (% of all TMP SAVs) | TMH Region | Non-TMH Region | Signal Peptide Region |
| --- | --- | --- | --- | --- |
| Common SAVs | 0.571 ± 0.006% | 0.62 ± 0.02% | 0.476 ± 0.003% | 1.10 ± 0.06% |
| Rare SAVs | 99.03 ± 0.07% | 99.0 ± 0.2% | 99.16 ± 0.01% | 98.2 ± 0.6% |

Looking at the variants in different TMP regions, the signal peptide region is strongly enriched in common variants and absent in rare variants by comparing it to the same proportion in all regions (0.571% and 99.03% respectively). The signal peptide region is, compared to the protein itself, very short, and its amino acid sequence is very sensitive to changes (a small change may drastically change the function of the signal peptide). It may be for this reason that only desirable variants are conserved, becoming more common. On the other hand, rare variants may be the result of undesired or non-functional changes for the signal peptide which have been discarded, becoming particularly less frequent.

Finally, non-TMH regions have a slightly higher proportion of rare variants and a lower one of common variants when compared to other regions. TMH regions, on the other hand, have a slight enrichment in common SAVs compared to all regions in general (0.62% vs 0.571%).

Next, we studied both the effect and frequency of variants in different regions (Table 10). Two main conclusions can be drawn. First, there is an enrichment of common, strong-effect SAVs in signal peptide regions, and a clear absence of their rare counterparts.

**Table 10.**
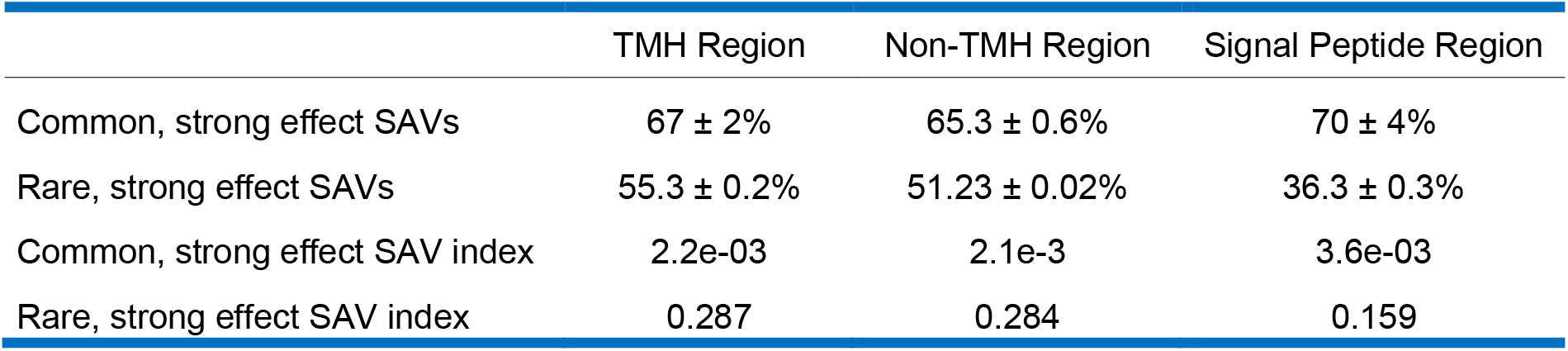
Distribution of common SAVs with a strong effect associated in the TMP regions. Last two rows show the variant / amino acid ratio (called index) to take into account the size of each region.

|  | TMH Region | Non-TMH Region | Signal Peptide Region |
| --- | --- | --- | --- |
| Common, strong effect SAVs | 67 ± 2% | 65.3 ± 0.6% | 70 ± 4% |
| Rare, strong effect SAVs | 55.3 ± 0.2% | 51.23 ± 0.02% | 36.3 ± 0.3% |
| Common, strong effect SAV index | 2.2e-03 | 2.1e-3 | 3.6e-03 |
| Rare, strong effect SAV index | 0.287 | 0.284 | 0.159 |

Even though the number of common, strong effect variants in the signal peptide regions is considerably small (only 230 variants meet this criteria), there can be a biological explanation for this enrichment. As mentioned earlier, this enrichment may also be the result of variants that had desired effects on protein function and were conserved, thus increasing their frequency to common levels. On the other hand, non-desired changes in the protein function will result in the variant being discarded, and its proportion decreasing.

This reasoning can easily be applied to all regions in TMPs: in all cases, the proportion of common variants with a strong effect associated is clearly higher than the proportion of rare variants (first two rows of Table 10). However, we can see how this effect is especially important in the signal peptide regions.

The second conclusion that can be drawn is that TMH regions have, proportionally, more common variants with a strong effect on protein function than non-TMH regions.

The enrichment found in TMHs could be due to several factors: on the one hand, the number of SAVs that meet these two requirements (common frequency, strong functional effect predicted) for this region is arguably small (852 variants meet this criteria). On the other hand, there may be a biological explanation for this enrichment: Since these regions have a specific composition that makes them suitable to go through the membrane (hydrophobic amino acids that favour a helical structure), the variety of SAVs that can affect the protein function without breaking the TMH structure is lesser.

Last of all, to complete the analysis of SAVs in different TMP regions, we studied the proportion of common and rare variants with a certain effect on protein function, grouped by the TMP region.

Regarding the SAV distribution in the TMP regions for different SNAP2 thresholds, we see that common variants, when grouped by the region where they belong, have proportionally a higher effect on protein function than rare variants (Fig. 1). This difference is visible in almost the complete threshold range (−100 to 100), and regardless of how the variants are divided: it can be observed when SAVs are divided by TMP types (Fig. 1), and by region (Fig. 3). In addition, TMH regions have a higher proportion of common variants with a strong effect associated compared to non-TMH regions.

**Figure 3.**
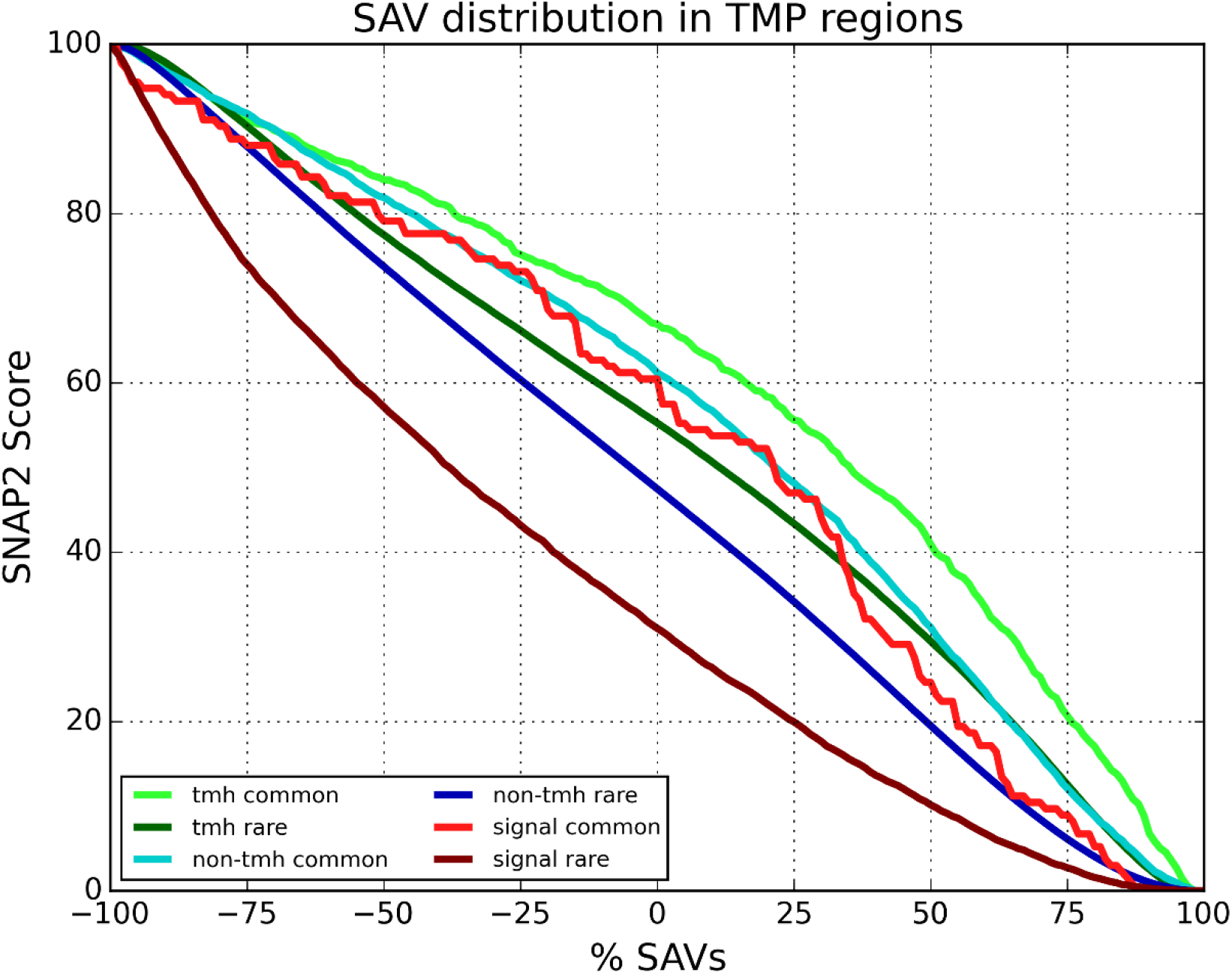
Common and rare variants, by region, predicted to have an effect on protein function. Variants have been grouped by the TMP region where they can be found. Y axis: Percentage of SAVs that have been predicted to have an effect. X axis: SNAP2 threshold used to predict that a variant has effect on protein function. Light colours stand for common variants, and darker colours for rare variants.

The line that belongs to common variants in signal peptides show a poor resolution, pointing out that there may be not enough common variants in these regions to draw a conclusion. Further analyses of these variants are therefore highly recommended, so the enrichment in common variants can be confirmed. The rare variants in the signal peptide regions show, on the other hand, a clear lack of effect on protein function. It confirms the idea that rare, undesirable effects on the signal peptides are discarded more easily, becoming particularly less frequent, therefore the rare variants that remain have progressively less effect on protein function.

## Conclusion

In this study we have analysed the single amino acid variants (SAVs) that can be found in the human proteome, focusing mainly on helical transmembrane proteins (TMPs). We have shown at the beginning, as a means of validation of our dataset, the present enrichment in common variants with a strong effect on protein function in all proteins of the human proteome.

Our first conclusion is that this SAV enrichment is also present in TMPs: all types of TMPs and all regions within TMPs show an enrichment in common variants with a strong effect on protein function. However, this enrichment is underrepresented compared to globular proteins.

The second conclusion, focused on TMP types, is that all types of TMPs show a clear enrichment in common, strong effect SAVs compared to rare variants. Regarding 7-pass TMPs in particular, there are slightly fewer variants in the TMH region, possibly due to the risk of modifying the complex structure of the protein, or its highly specific function. Moreover, there were no common variants with a strong effect associated in the signal peptides from 7-pass proteins, and there were only 20 of their rare counterparts, which can be interesting to look further into them.

Our third conclusion covers the TMP regions. TMH regions are more enriched in common variants with a strong effect on protein function for almost all SNAP2 thresholds. Signal peptides have a clear reduction of rare variants with a strong effect associated. Although there are not many variants that meet these criteria in signal peptide regions, the reason for this enrichment may be biological: the modified signal peptide may have a relevant function as well, and it may be of interest to keep, therefore increasing its frequency.

In general, there is a higher proportion of common, strong effect SAVs in all TMP types and regions compared to their rare counterparts. This difference is observable using any SNPA2 threshold to determine the strength effect of the variant, especially with non-extreme threshold values (Fig. 1-3).

Lastly, it remains to perform a more thorough analysis of several interesting groups of variants: those from very small groups that did not allow to draw a definitive conclusion (i.e. strong effect variants in signal peptide regions of 7-pass TMPs), and variants for the most relevant enrichments (i.e. common variants in TMH regions, and common and rare variants in signal peptide regions). Said enrichments may be better explained once we analyse the specific change in protein function of the variants, the exact location of the variants, whether a specific location for the variants may bring similar changes in the protein function, etc. We confirm that signal peptides, normal TMH regions, and TMPs in general, are enriched in common variants with a strong effect on protein function.

## Abbreviations used

GPCR: G protein-coupled receptors (7-pass TMPs)
non-TMP: non-transmembrane proteins (due to dedication errors, these may include some beta-barrel membrane proteins)
SAV: single amino acid variant
SNV: single nucleotide variant
TMH: transmembrane helix
TMP: transmembrane protein.

## Acknowledgements

Thanks to the authors who made available the tools that were used here, and those who also deposit their data in public databases for further use. Special thanks to Tim Karl and Inga Weise (administration from TUM) for their assistance.

